# On the functional annotation of open-channel structures in the glycine receptor

**DOI:** 10.1101/2020.05.05.076802

**Authors:** Adrien Henri Cerdan, Marco Cecchini

## Abstract

The glycine receptor (GlyR) is by far the best-characterized pentameric ligand-gated ion channel with several high-resolution structures from X-ray crystallography, cryo-EM and modeling. Nonetheless, the significance of the currently available open-pore conformations is debated due to their diversity in the pore geometry. Here, we discuss the physiological significance of existing models of the GlyR active state based on conductance and selectivity measurements by computational electrophysiology. The results support the conclusion that the original cryo-EM reconstruction of the active state obtained in detergents as well as its subsequent refinement by Molecular Dynamics simulations are likely to be non-physiological as they feature artificially dilated ion pores. In addition, the calculations indicate that a physiologically relevant open pore configuration should be constricted within a radius of 2.5 and 2.8Å, which is consistent with previous modeling, electrophysiology measurements, and the most recent cryo-EM structures obtained in a native lipid-membrane environment.

---

The functional annotation of the high-resolution structures of biological ion channels from X-ray crystallography and cryo-electron microscopy is a difficult task, which is crucial for a molecular understanding of synaptic transduction and its chemical regulation by pharmacological agents (Cecchini and Changeux, 2015). The glycine rceptor (GlyR) is by far the best characterized pentameric channel with several high-resolution structures solved in different conformational states and in complex with modulatory ligands, i.e., agonists, antagonists, partial agonists, and allosteric modulators. Perhaps surprisingly, two apparently different structures from cryo-EM could be associated with an open state, i.e. the *wide-open* structure featuring an ion pore as large as 4.4 Å in radius, and the significantly more contracted *semi-open* structure with a pore radius of 2.4 Å at the constriction point (Du et al., 2015). Functional annotation based on all-atom Molecular Dynamics (MD) challenged both the *wide-open* structure, whose pore is too wide to be blocked by picrotoxin (Gonzalez-Gutierrez et al., 2017) and is too conductive and non-selective to the size of the anion (Cerdan et al., 2018), and the *semi-open* structure, whose pore appeared as intermittently dehydrated in MD (Trick et al., 2016) and non-permeable to chloride at physiological conditions (Cerdan et al., 2018). Furthermore, an alternative conformation of the receptor with an intermediate pore radius (2.5 Å), which was recently captured by MD (*MD-open*), was proposed to provide a better representation of the physiologically active state (Cerdan et al., 2018).

In a recent contribution in *Structure* (Dämgen and Biggin, 2019), Dämgen and Biggin report on a fourth open-channel structure of GlyR, here *D&B-open*, which features an ion pore halfway between the *wide-open* and *MD-open* states with a radius of 3.6 Å. The enhanced structural stability of *D&B-open* relative to the parental *wide-open* structure was attributed to the occupancy of a large hydrophobic cavity in the trans-membrane domain at the interface between subunits by an alternative rotameric state of the pore-lining leucines at position 9′. Based on the structural stability in MD and both hydration and chloride permeability of the ion pore, these authors concluded that *D&B-open* is functionally open and provides the best model for the active state.

Using computational electrophysiology on a reduced molecular system composed of the trans-membrane domain only in a lipid bilayer and harmonic restraints on the heavy atoms (Figure 1), we have analyzed the ion conductance and selectivity of *D&B-open* and compared with single-channel electrophysiology; see SI for a justification of the setup. The numerical results collected at a transmembrane potential of 150*mV* or 250*mV*, which was introduced via a constant electric field (Roux, 2008; Gumbart et al., 2012) or charge imbalance (Sachs et al., 2004; Kutzner et al., 2011) (Figure S1 and SI), indicate that the conductance of *D&B-open* ranges from 382*ps* to 480*ps*, which is about five-fold higher than the experimental value of 86*pS* (Bormann et al., 1993; Moorhouse et al., 2002; Moroni et al., 2011a; Scott et al., 2015; Lara et al., 2019); see Table S1. Moreover, *in-silico* permeation assays using a series of polyatomic anions, which were originally used to probe the size of the GlyR ion pore at physiological conditions (Bormann et al., 1987; Rundstrom et al., 1994; Lee et al., 2003), indicate that *D&B-open* is not only too conductive but also non-selective to the size of the anion, in clear contradiction with the experiments; see Table S2. Taken together, these numerical results support the conclusion that *D&B-open* is non-physiological as much as the parental *wide-open* structure, despite a significant reorganization of the pore-lining residues that result in a narrower ion pore. Remarkably, this prediction appears to be supported by the most recent cryo-EM reconstructions of the GlyR channel isolated in complex with endogenous lipids (Yu et al., 2019). In fact and unlike *D&B-open*, the open channel with glycine bound features a pore radius of 2.8 Å, in agreement with previous electrophysiological recordings with polyatomic anions (Bormann et al., 1987; Rundstrom et al., 1994; Lee et al., 2003) (2.65 – 2.8 Å) and simulation results (*MD-open*, 2.5 Å) (Cerdan et al., 2018), see Table 1. Moreover, in the new cryo-EM open state, the pore-lining helices M2 are tilted solely in the polar direction consistent with *MD-open* and in contradiction with the significant azimuthal tilting observed in the *wide-open* and the *D&B-open* structures. Last, concerning the role of the pore-lining leucines at position 9′ on the stabilization of the physiological active state of GlyR, which was raised by Dämgen and Biggin (Dämgen and Biggin, 2019), the debate seems not to be settled yet. In fact, the very recent cryo-EM structures of GlyR-*α*1 in native lipid environments indicate that these leucines undergo a more pronounced rotation (away from the lumen) in the desensitized state rather than the active state. But, this conclusion is not supported by the very recent cryo-EM analysis of *Kumar et al.* (Kumar et al., 2019).

**Figure 1:**
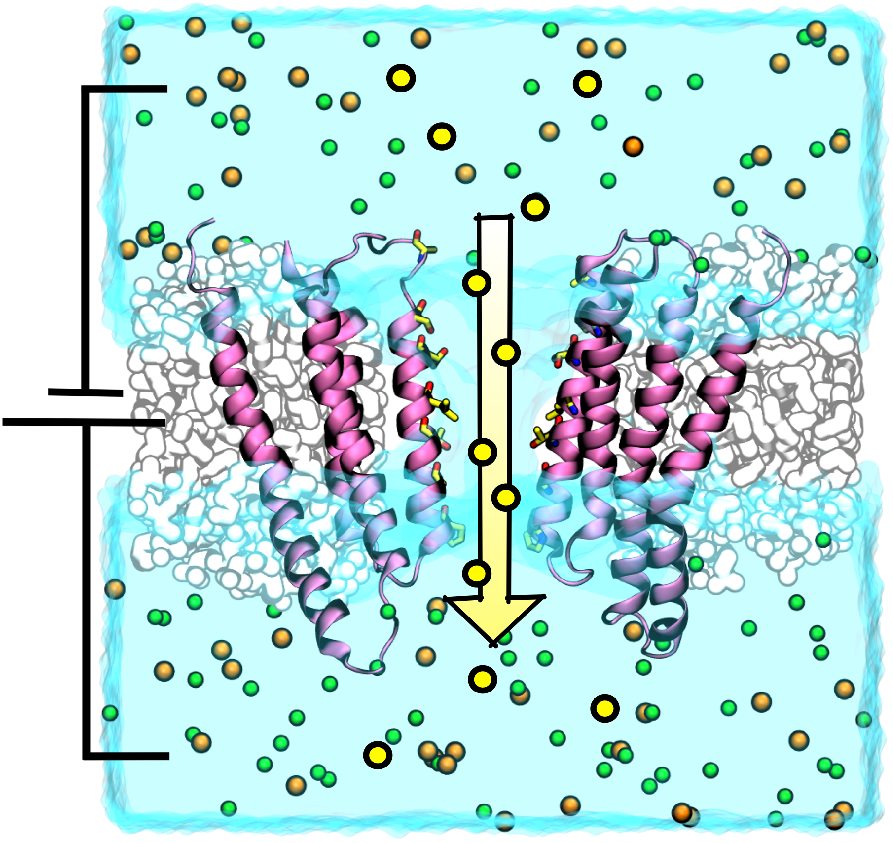
Computational electrophysiology of the GlyR channel by all-atom Molecular Dynamics. The ion-channel is shown in violet, the lipid membrane in white, and sodium and chloride ions are represented as green and yellow balls, respectively. In this setup, a truncated construct (TMD-only) embedded in a lipid bi-layer and exposed to a physiological concentration of chloride is simulated. A physiological membrane potential is introduced using a constant electric field perpendicular to the membrane. Channel conductance is measured from the ionic flux (yellow arrow) across the membrane divided by the trans-membrane voltage. The removal of the extracellular domain (ECD) reduces the system’s size to approximately one half; compare ~95, 000 (reduced) with ~200, 000 (original) atoms. See SI for details.

**Table 1:**
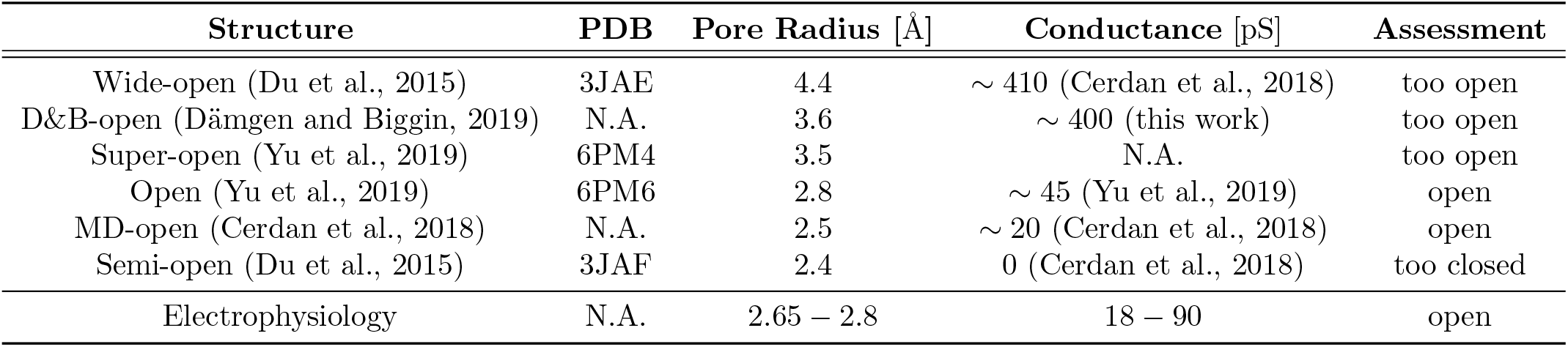
Summary of existing *open-state* structures of GlyR-*α*1 with an assessment on their relevance based on computational electrophysiology. The experimental conductance values were obtained from single-channel electrophysiological recordings (Bormann et al., 1993; Moorhouse et al., 2002; Moroni et al., 2011a; Scott et al., 2015; Lara et al., 2019), and the minimal pore radius was estimated from permeability assays to polyatomic anions (Bormann et al., 1987; Rundstrom et al., 1994; Lee et al., 2003). See also Table S1 and Table S3 for details.

The observations above lead to the following conclusions. First, computational electrophysiology results along with cryo-EM reconstructions in native lipid environments strongly support the physiological relevance of the openchannel structure that we have recently isolated by Molecular Dynamics (Cerdan et al., 2018); see Table 1. Second, our analysis of *D&B-open* indicates that structural stability and water permeation in MD is necessary but not sufficient to assess the relevance of an open-channel structure. Third, computational electrophysiology in conjunction with polyatomic anion permeability, whose implementation is straightforward in modern MD engines, provides useful information for the functional annotation, which helps to discriminate between alternative open-pore structures. Fourth, both modeling (Cerdan et al., 2018) and structural biology (Yu et al., 2019) consistently indicate that the GlyR conductance at physiological conditions, i.e. 18 88 pS (Bormann et al., 1993; Moorhouse et al., 2002; Moroni et al., 2011a; Scott et al., 2015; Lara et al., 2019), does not require the permeation of fully hydrated chloride ions (i.e. 3.2 Å in radius). Based on these observations, we speculate that partial dehydration might possibly provide the best compromise between conductance and selectivity in GlyR and possibly other anionic channels. Although ion conductance and selectivity predicted by MD are not expected to be in quantitative agreement with the experiments (i.e. the calculated values are subjected to systematic errors from the force field, which dictate the ion diffusion coefficient and the strength of the protein-ion interaction within the ion pore), these calculations provide at least order-of-magnitude estimates that may be used to rule out non-physiological states. In this respect, the implementation of computational electrophysiology based on polarizable force fields appears as a promising tool by which to interrogate ion channel structures more quantitatively and establish relationships between structural and functional states (Klesse, 2019).

## Supporting Information

### Detailed Results

#### Conductance

To evaluate the physiological significance of the *D&B-open* structure, its ion conductance, and selectivity were analyzed by computational electrophysiology based on all-atom MD and compared to single-channel electrophysiology results. By applying a constant external electric field that emulates a trans-membrane potential of 150*mV* and 250*mV* (Roux, 2008; Gumbart et al., 2012), we recorded conductance values of 395 and 426*pS*, respectively, during 200*ns* simulations (see Table S1). The numerical results indicate a conductance that is almost five-fold higher than the experimental value of 86*pS*. Remarkably, the computed conductance for *D&B-open* is similar to the one measured in the non-physiological *wide-open* conformation (413*pS*) (Cerdan et al., 2018), despite the pore radius at the constriction point is 0.8 Å narrower. This observation suggests that the *D&B-open* conformation, which corresponds to a structurally stable alternative of the *wide-open* state, is equally non-physiological.

To verify that the constant electric field does not bias the ion translocation process and therefore the measured conductance, we also tested the permeability of *D&B-open* within the double membrane ion-imbalance setup (Kutzner et al., 2011) which emulates the physiological conditions more closely. By imposing a charge-imbalance (∆*q*) of 4*e* or 6*e*, the resulting transmembrane potential amounts to 180 mV and 270 mV, respectively, and the computed conductances 383 and 481*pS* over 200*ns* simulations (see Table S1). Overall, conductance values computed by both methods are similar enough to conclude that the application of an external electric field of 0.014 or 0.024 V/nm does not bias the chloride translocation with respect to the more *natural* charge-imbalance protocol.

#### Polyatomic Anion selectivity

When applied to *D&B-open* (see Table S2), our simulation protocol highlight a similar pattern to the one observed in the *wide-open* conformation, i.e., the whole range of tested anions is probed to be permeable.

### Materials and Methods

#### System preparation

The initial coordinates of the *D&B-open* structure of the human GlyR-*α*1 were obtained from *Dämgen and Biggin, 2019* (Dämgen and Biggin, 2019) (DOI: 10.5281/zenodo.3476169). Following the protocol in *Cerdan et al., 2018* (Cerdan et al., 2018), the extracellular domain was removed and the transmembrane domain was embedded in a POPC membrane bilayer, surrounded by TIP3 water molecules and 150 mN NaCl using the CHARMM-GUI input generator (Lee et al., 2016, 2019). The simulation box contains 148, 106 atoms, including one protein, 340 POPC molecules, 30, 599 water molecules, 82 Na^+^, and 122 Cl^−^ The initial box dimensions are 120 ∗ 120 ∗ 111Å.

To generate a dual membrane setup, we adapted the script *makeSandwich.sh* provided by *Kutzner et al., 2011* (Kutzner et al., 2011) (available at https://www.mpibpc.mpg.de/grubmueller/compel) to the latest GROMACS version (2019.4) (Abraham et al., 2015).

To evaluate polyatomic-anions relative permeabilities, we used parameters from *Cerdan et al., 2018* (Cerdan et al., 2018), which were produced using CGenFF through the PARACHEM website (Vanommeslaeghe et al., 2010) (https://cgenff.paramchem.org). For the polyatomic-anions experiment, we *ionized* the system at 1 M NaCl using VMD (Humphrey et al., 1996) and subsequently replaced the chloride atoms by polyatomic-anions removing the overlapping water molecules.

#### MD Simulations

All MD simulations were run with GROMACS 2019.4 (Abraham et al., 2015), using the periodic boundaries conditions, and the CHARMM36 forcefield (Best et al., 2012; Klauda et al., 2010) with CHARMM36m (Huang et al., 2017) modifications. The minimization and equilibration protocols used default parameters generated by CHARMM-GUI for GROMACS. In short, after 5, 000 steps of minimization using the steepest-descent algorithm, the system was heated at 300 K by generating random velocities with the Berendsen thermostat for 50 ps using a 1 fs integration timestep. Then, the system was coupled semi-isotropically to a Berendsen barostat and further equilibrated for 25 ps using 1 fs timestep, and 200 ps with a 2 fs timestep. During the equilibration, atomic positions restraints on protein heavy atoms were gradually relaxed from 4, 000 and 2.000 kJ mol^−1^ nm^−2^ to 500 and 200 kJ mol^−1^ nm^−2^ for protein backbone and side-chains atoms, respectively. The production runs were carried out using a 2 fs timestep in the NVT ensemble, with velocity-rescaling to maintain a temperature of 300 K. Atomic positions restraints were maintained on protein heavy atoms with force constants of 500 and 200 kJ mol^−1^ nm^−2^ on the backbone and side-chains atoms, respectively. Generally, the LINCS algorithm was used to constraints the H-bounds and the Particle Mesh Ewald (PME) to treat the long-range electrostatics.

#### Computational Electrophysiology

To evaluate channel conductance *in silico*, two computational electrophysiology setups were used. The first relies on the introduction of a constant electric field (*E_z_*) along the direction perpendicular to the membrane plane (*z*). The resulting transmembrane potential (*V_m_*) is proportional to the applied electric field and the length of the simulated box 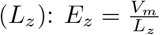 (Roux, 2008). Based on the size of the simulations box at the end of the equilibration, electric fields of 0.014 or 0.024 V/nm were applied along the *z*-axis to introduce transmembrane potentials of 150 and 250 mV, respectively.

Estimates of the ionic currents were obtained by counting the number of permeation events per unit of time using the FLUX module in Wordom (Seeber et al., 2007, 2011). Error estimates (*σ*) on the current (*I*) are provided assuming a Poisson distribution of the permeation events such that 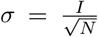, with *N* the number of events (Sotomayor et al., 2007). Finally, a channel conductance (*g*) was computed as 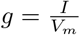.

**Table S1:** Conductance of the *D&B-open* conformation computed *in silico* by all-atom MD at 150mM NaCl. Two methods were used to generate the transmembrane potential, an external Electric Field (EF) or a Charge-Imbalance (CI). The constant electric field applied on all atoms is directly proportional to the transmembrane potential and the size of the simulated box along the z-axis (normal to the membrane plane). To produce comparable voltage-drops across the membrane we introduced an imbalance of 4 or 6 charges. The selectivity for chloride over sodium could not have been quantitatively evaluated because no permeable sodium have been recorded during the simulation time, indicating a high selectivity for chloride.

| Setup | Permeation Events | Current (pA) | Conductance (pS) | Selectivity ($P_{Cl^-}/P_{Na^+}$) | Total simulation time (nS) |
| --- | --- | --- | --- | --- | --- |
| EF: 150mV (replica 1) | 39 | $62.5 \pm 10.0$ | $416.6 \pm 66.7$ | N.D. | 100 |
| EF: 150mV (replica 2) | 35 | $56.1 \pm 9.5$ | $373.8 \pm 63.3$ | N.D. | 100 |
| EF: 150mV (total) | 74 | $59.3 \pm 6.9$ | $395.2 \pm 45.9$ | N.D. | 200 |
| EF: 250mV (replica 1) | 70 | $112.2 \pm 13.4$ | $448.6 \pm 53.6$ | N.D. | 100 |
| EF: 250mV (replica 2) | 63 | $100.9 \pm 12.7$ | $403.8 \pm 50.8$ | N.D. | 100 |
| EF: 250mV (total) | 133 | $106.5 \pm 9.2$ | $426.2 \pm 37.0$ | N.D. | 200 |
| CI: 180mV | 86 | $68.9 \pm 7.4$ | $382.7 \pm 41.3$ | N.D. | 200 |
| CI: 270mV | 162 | $129.8 \pm 10.2$ | $480.7 \pm 37.8$ | N.D. | 200 |

**Table S2:** Permeability Ratio 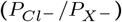 of Polyatomic Anions in the Open Structures of GlyR-*α*1. The results obtained for *wide-open* and *MD-open* conformations come from (Cerdan et al., 2018), the experimental values from (Bormann et al., 1987), and the data concerning *D&B-open* corresponds to the present contribution. The first four anions are experimentally permeable through the pore of GlyR-*α*1, while the four last are not permeable. The external electric field method was used in this experiment in order to introduce a transmenbrane potential corresponding to 150 mV with 1 M ionic concentration.

| Anions | *wide-open* | *D&B-open* | *MD-open* | Exp. |
| --- | --- | --- | --- | --- |
| Chloride | **1** | **1** | **1** | **1** |
| Formate | **0.65** | **0.58** | **1.39** | **0.33** |
| Bicarbonate | **0.37** | **0.32** | **0.17** | **0.11** |
| Acetate | **0.32** | **0.38** | **0.18** | **0.035** |
| Phosphate | **0.15** | **0.22** | **0** | **0** |
| Methanesulfonate | **0.38** | **0.29** | **0** | **0** |
| Propionate | **0.24** | **0.21** | **0** | **0** |
| Isethionate | **0.19** | **0.19** | **0** | **0** |

**Figure S1:**
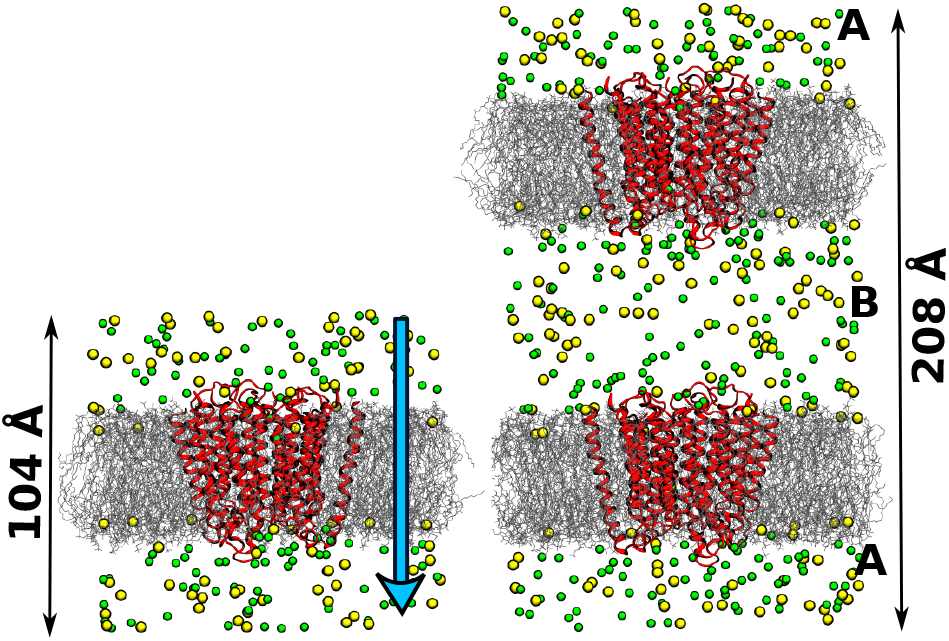
MD setup for computational electrophysiology. On the left, the *constant electric field* setup is shown. In this case, an external electric field along the z-axis (blue arrow) is applied, which is proportional to the transmembrane potential and the simulated box length. On the right, the *charge imbalance* setup is shown. In this case, two distinct compartments (A and B) are simulated, which allows for the introduction and conservation of a charge-imbalance, ultimately generating the desired transmembrane potential. Both setups rely on the use of the *reduced* molecular system containing only the TMD embedded in a membrane bilayer.

Concerning the charge-imbalance method (Kutzner et al., 2011), channel conductance was evaluated using the *CompEL* protocol as implemented in GROMACS 2019.4 with default parameters. Upon the introduction of the charge imbalance (∆*q*), the resulting transmembrane potential was measured using the *gmx potential* program. The transmembrane potential produced by a ∆*q* of 4*e* or 6*e* corresponds to 180 or 270 mV, respectively. Permeation events were extracted from the default output *swapions.xvg* in order to compute currents, conductances, and corresponding errors as described above.

#### On the use of a truncated model of GlyR

It has been argued that the use of a truncated model of GlyR, i.e. a model devoid of both ECD and ICD, may provide calculated conductance values that are not directly comparable with the experimental measurements, particularly if the rate-limiting step for ion translocation is not located in the TMD (*Biggin et al.*, this issue). Although there exist several pieces of evidence that support the role of the ECD and ICD on the modulation of ion conductance in pLGICs, we argue that these observations were mostly done with cationic channels and they do not necessarily apply to GlyR. In fact, our working hypothesis that the rate-limiting step for the chloride translocation in homomeric GlyR-*α*1 is located in the TMD is supported by the following evidence:

1. All mutations modulating conductance in GlyR that were identified in the ECD have a negative effect on the conductance, i.e., they reduce conductance by introducing a barrier rather than removing it (Moroni et al., 2011b; Brams et al., 2011).
2. Unlike cationic channels, all mutations modulating conductance in GlyR that were identified in the ICD have a negative effect on the conductance, i.e., they reduce conductance by introducing a barrier rather than removing it (Carland et al., 2009).
3. Homomeric GlyR-*α*1 in absence of the ICD (truncated M2-M3) displays the same conductance as the full-length wild type receptor (Moroni et al., 2011a,b).
4. The chimeric receptor Lily, which was obtained by fusing the ECD of GLIC and with the TMD of GlyR in absence of ICD (truncated M3-M4 loop), named displays the same conductance as the fulllength wild type GlyR-*α*1, including the existence of sub-conductive states (Moraga-Cid et al., 2015).
5. To the best of our knowledge, the only mutation that increases the single-channel conductance in GlyR a1 is located in the 2’ position of the TMD (G221A). The effect of this mutation is produced both in full-length wild-type (Bormann et al., 1993) and the chimeric receptor Lily (Moraga-Cid et al., 2015).

Therefore, the existing literature on GlyR-*α*1 is consistent with the conclusion that the rate-limiting step for chloride translocation is located in the TMD, which justifies the use of a truncated TMD-only model to evaluate the conductance by computational electrophysiology.

#### On the use of position restraints

It has been argued that the use of position restraints on the protein backbone, which are required to explore the conductance and selectivity of a truncated form of an ion channel, may prevent correct sampling of the true conformational ensemble of the physiological active state (*Biggin et al.*, this issue). Since we agree that fully unrestrained simulations, albeit computationally more intensive, would provide a better representation of the ion-channel conductance and selectivity, the full-length cryo-EM construct of the GlyR *α*1 (ECD+TMD) was explored by computational electrophysiology starting with the structure of the MD-open state (Cerdan et al., 2018). Upon running multiple repeats (10 per voltage) of long (200 – 300*ns*) and fully unrestrained simulations in the presence of a membrane potential of 150*mV* or 250*mV*, we observed a slight increase in the conductance (from ~20*pS* to ~30*pS*) relative to the truncated form in the presence of position restraints; see Table S3. Intriguingly, the release of the position restraints appears to increase channel conductance by promoting larger fluctuations at the level of the ion pore. These results demonstrate that the use of position restraints has only a minor effect on the channel conductance, which supports the assessment on the physiological relevance of the various GlyR open-state structures based on computational electrophysiology of truncated constructs; see Table 1.

**Table S3:**
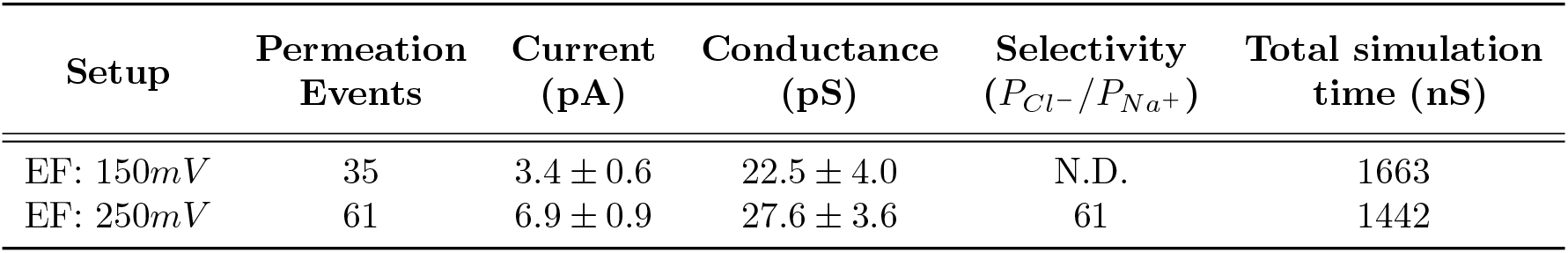
Calculated conductance of the “full-length” (ECD+TMD) *MD-open* state by all-atom MD at 150mM NaCl in the absence of position restraints on the protein backbone. Permeation events were counted over simulated trajectories collected in the presence of an external electric field. For each voltage, multiple replica simulations were carried out for 200 – 300*ns*. Statistics were computed by averaging permeation events over all replicas at each voltage.

| Setup | Permeation<br>Events | Current<br>(pA) | Conductance<br>(pS) | Selectivity<br>($P_{Cl^-}/P_{Na^+}$) | Total simulation<br>time (nS) |
| --- | --- | --- | --- | --- | --- |
| EF: 150mV | 35 | $3.4 \pm 0.6$ | $22.5 \pm 4.0$ | N.D. | 1663 |
| EF: 250mV | 61 | $6.9 \pm 0.9$ | $27.6 \pm 3.6$ | 61 | 1442 |

